# The H3K27me3 epigenetic mark is crucial for callus cell identity and for the acquisition of new fate during root and shoot regeneration

**DOI:** 10.1101/2022.05.12.491615

**Authors:** Tali Mandel, Udi Landau, Tommy Kaplan, Leor Eshed Williams

## Abstract

We combined molecular, genomic and genetic approaches to study the molecular mechanisms underlying cell totipotency and competency to regenerate in Arabidopsis.

By performing comparative analysis of mRNA-seq and chromatin landscapes between leaf differentiated cells and callus totipotent cells and between WT callus and calli derived from the *emf2* mutant, exhibiting impaired regenerative capacity we revealed the following:

**1**. That callus cells express numerous genes of many developmental pathways such as root, leaf, embryo, shoot, meristem and seed. This suggests a mechanism to allow rapid response to a signal by maintaining genes of all potential developmental pathways active, without the needs to release transcriptional silencing and to go through the intricate multistep process of transcription. **2**. That key transcription factors that are sufficient to derive differentiation or organogenesis are silenced and marked by the H3K27me3. **3**. That callus derived from the *emf2* mutant which is impaired in setting the H3K27methylation, lost the capacity to regenerate and that 78 transcription factors from which 18 regulate flower development, where up-regulated compared with WT callus.

Altogether our results suggest that competency to regenerate is achieved by keeping the chromatin of developmental genes active, and that upon a signal for cell fate switch, a mechanism to repress those genes is required to allow the one desired developmental pathway to dominate. When this mechanism is impaired the capacity to regenerate is decline.

## Introduction

Certain plant cells retain plasticity and are capable - under appropriate stimuli, e.g. after wounding or following hormone stimulation - to switch their developmental program, re-enter the cell cycle and form a mass of less-differentiated pluripotent cells, termed callus (1). Those cells acquire competence to respond to diverse stimuli, taking on new fate accordingly. For example, high auxin/cytokinin ratio induces root regeneration, whereas a low ratio promotes shoot induction (2). In multicellular organisms the acquisition of new cell fate and maintenance of cellular identity require the establishment of specific gene expression patterns and a cellular memory system to transfer these patterns during cell divisions.

Tri-methylation of histone H3 Lys27 (H3K27me3) which is catalyzed by the Polycomb Repressive Complex 2 (PRC2), serves as an epigenetic repressive chromatin mark, central to the control of developmental genes expression during cell fate transitions and lineage specification. The function of PRC2 and its components was originally discovered in *Drosophila melanogaster*, in mutants displaying homeotic conversions due to ectopic expression of homeotic (Hox) genes (3). The PRC2 is composed of four components, highly conserved across higher organisms (4), and it is the sole known complex with H3K27me3 methyltransferase activity.

The importance of PRC2 and H3K27me3 in numerous organisms is highlighted by the severe developmental defects and even lethality, caused by mutation or depletion of PRC2 subunits (5). The four core subunits of the Drosophila PRC2, each encoded by a single copy gene, are the catalytic component Enhancer of zeste (E(z)) which has the signature SET domain; the non-catalytic partners, Suppressor of zeste 12 (Su(z)12) and Extra sex combs (Esc), which directly contact E(Z) and are indispensable for the methyltransferase activity; and the Nucleosome remodeling factor 55 kDa (Nurf55).

In the flowering plant Arabidopsis thaliana, PRC2 core subunits, apart from the Esc homologue FERTILISATION INDEPENDENT ENDOSPERM (FIE) are encoded by multigene families. The core subunits are assembled in different combinations into three distinct PRC2 complexes name after the Su(z)12 homologs component: the EMBRYONIC FLOWER2 (EMF2), the FERTILISATION INDEPENDENT SEED (FIS2), and the VERNALISATION 2 (VRN2) (6). Each complex controlling divers process and developmental programs with some redundancy, including cell fate specification, embryogenesis, vegetative growth and flowering time.

In plant, the genomic patterns of H3K27me3 is highly dynamic and strongly associated with transcriptional repression of hundreds of genes many which are developmental. Massive reprograming of H3K27me3 is taking place during the acquisition of pluripotent state at the formation of callus (7) and during differentiation of stem cell at the shoot apical meristem (8).

## Results

### Callus pluripotent cells express numerous genes from many developmental pathways

To facilitate our understanding of pluripotency and competency to regenerate we compared the transcriptomes of Arabidopsis six-week old calli derived from cotyledons callus, which exhibits high capacity to regenerate, with leaves of three-week old plant, that in Arabidopsis show little or no capacity for direct regeneration (Supplementary Fig. 1) (9).

In total 18,617 genes were found to be expressed (RPKM>1) in all samples, 16,703 genes in callus and 16,694 in leaves (Supplementary Data 1 and 2), with 12,109 genes exhibiting significant differential expression (p – Value <0.5) (Fig. 1a and Supplementary Data 3), demonstrating the substantial dissimilarity between cells comprising the callus and the leaf. As expected, Gene Ontology (GO) analysis showed that the 6223 callus down-regulated genes (i.e. up-regulated in leaves) are significantly enriched for genes related to photosynthesis (p<E-76) in the biological processes term, and plastid and thylakoid (p<E-69) in the cellular component terms (Supplementary Data 4). This reflects the function of the leaf as photosynthesis factories and indicates that the leaf cells are differentiated (10). The GO enrichment for the 5886 callus up-regulated genes is shown in Figure 1b (full terms list in Supplementary Data 5). Out of the 3425 genes in the nucleic acid metabolic term (GO:0090304), 2574 are expressed in callus (p<E-75), suggesting high production of nucleic acids that are required for DNA synthesis in the highly proliferative cells, consistent with the enrichment in cell cycle genes (316 are expressed out of 423 in the term (GO:0007049)). The enrichment in response to endogenous stimulus (GO:0009719) was also predictable as the callus is cultured on media supplemented with the phytohormones auxin and cytokinin. However, what caught our attention is the significant enrichment in multicellular organismal development genes (GO:0007275). This was unexpected since callus consider to be an unorganized less differentiated mass of cells, and because it was shown by many to be enriched for root developmental genes (11, 12). A detailed GO analysis on the 1950 callus expressed genes that fall within the term (Supplementary Data 6), confirmed that 339 genes are related to root development (GO:0048364; 432 genes), including the well-established root marker genes, *WUSCHEL RELATED HOMEOBOX 5* (*WOX5*), *SHORT ROOT* (*SHR*) and *SOMBRERO* (13–15). But most striking was the significant enrichment of genes participating in many other developmental pathways including shoot, leaf, flower, embryo and seed development (Fig. 1c). For example, out of the 282 genes in the leaf development term (GO:0048366), 197 (70%) are expressed in callus, including key transcription factors (TFs) essential for leaf morphogenesis such as *ASYMMETRIC LEAVES 1* (*AS1*), *CUP-SHAPED COTYLEDON 2* (*CUC2*), *AUXIN RESPONSE FACTOR 3*/and *4* (*ARF3/4*) and the five members of the Class *III HD-ZIP* genes (16). Another example is the enrichment in meristem development (135 out of 174 in the term - GO:0048507), from which many genes are participating in shoot and inflorescent meristem development, like CORYNE, *KNOTTED1-LIKE HOMEOBOX GENE 6* (*KNAT6*) and *KNAT1* (17). Remarkably, genes known to be expressed specifically in the pluripotent stem cells at the SAM, like the *CLAVATA3* (*CLV3*), *AINTEGUMENTA-LIKE 7* and *INHIBITOR OF GROWTH 1* (18, 19), are not expressed in the callus.

**Figure 1:**
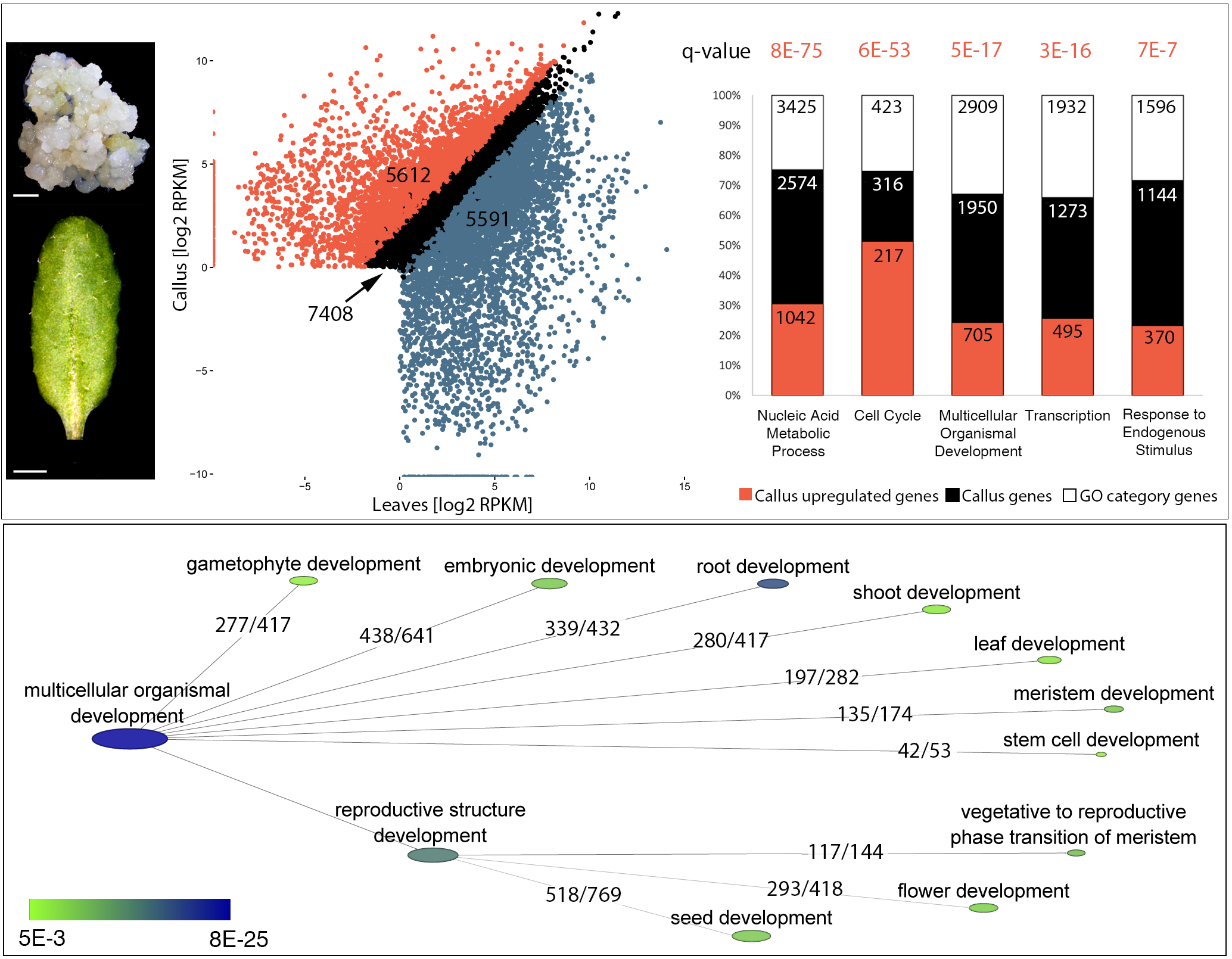
WT callus exhibits enrichment in genes participating in many developmental pathways. Five week-old cotyledon derived callus and 18-day old leaves were subjected to mRNA-seq analysis in 3 replicates. (**a**) Differential analysis reveals over 11,000 differentially expressed genes (callus significant up- and down-regulated genes are marked in red or blue respectively). **(b)** The enriched GO terms of biological functions for the callus expressed genes. The number of genes in the term (white), the callus expressed genes (black) and the callus significant up-regulated gene as compared with leaves (red). Nucleic acid metabolic term (GO:0090304), Cell cycle (GO:0007049), multicellular organismal development genes (GO:0007275), transcription (GO:0006350), response to endogenous stimulus (GO: 0009719). **(c)** Detailed GO analysis for the 1950 callus expressed genes of the multicellular organismal development gene term. For each pathway the number of genes expressed in the callus/out of the genes in the term are indicated. The full list of genes in each term is found in supplementary data X.

The expression of numerous genes from diverse developmental programs in the callus might provide means to respond rapidly to stimuli and to coordinate *de novo* organogenesis without the needs to release transcriptional silencing and to go through the intricate multistep process of transcription. In this scenario, we hypothesized that genes encoding for TFs that their expression alone is sufficient to direct differentiation or *de novo* organogenesis, must be silenced to confer pluripotency and to maintain callus cell identity. Indeed, genes which were shown to induce cell differentiation into stomata, trichrome, xylem fiber and tracheary cells, root hair and others, are not expressed (RPKM<1) in the callus (examples are given in Supplementary Data 7). The SPEECHLESS (SPCH), MUTE and FAMA TFs regulating stomatal development by promoting asymmetric divisions, acquisition of guard mother cell identity and guard cell differentiation respectively (20, 21), are one example. Key TFs that are necessary and sufficient to trigger *de novo* organogenesis, for instance the *WUSCHEL* (*WUS*) gene, that its expression alone in callus, leaves or roots, induces shoot meristem formation (22, 23), the *LEAFY* (*LFY*) that can direct flower formation (24) or *AGAMOUS* (*AG*), that its miss-expression leads to carpel formation (25, 26) are also not expressed in the callus.

### Enrichment of H3K4me3 in callus correlates with gene expression

Callus cells have high capacity to respond to diverse stimuli and to differentiate accordingly (27). The acquisition of new fate and maintenance of cell identity requires the establishment of lineage-specific transcriptional programs mediated primarily by epigenetic mechanisms (28). To characterize the H3K4me3 and H3K27me3 epigenetic marks, known to be associated with transcriptional activation and repression, respectively (29, 30), we performed ChIP-seq analyses on nuclei isolated from callus, using H3K4me3 and H3K27me3 specific antibodies. In total 16,161 genes were found to be marked by H3K4me3 (Supplementary Data 8) with an enrichment typically covering ~800bp of the gene, starting at the transcription start site (TSS) and peaking within ~ 200bp downstream to the TSS (Fig. 2a), consistent with the hallmark of H3K4me3 at gene TSSs (30).

**Figure 2.**
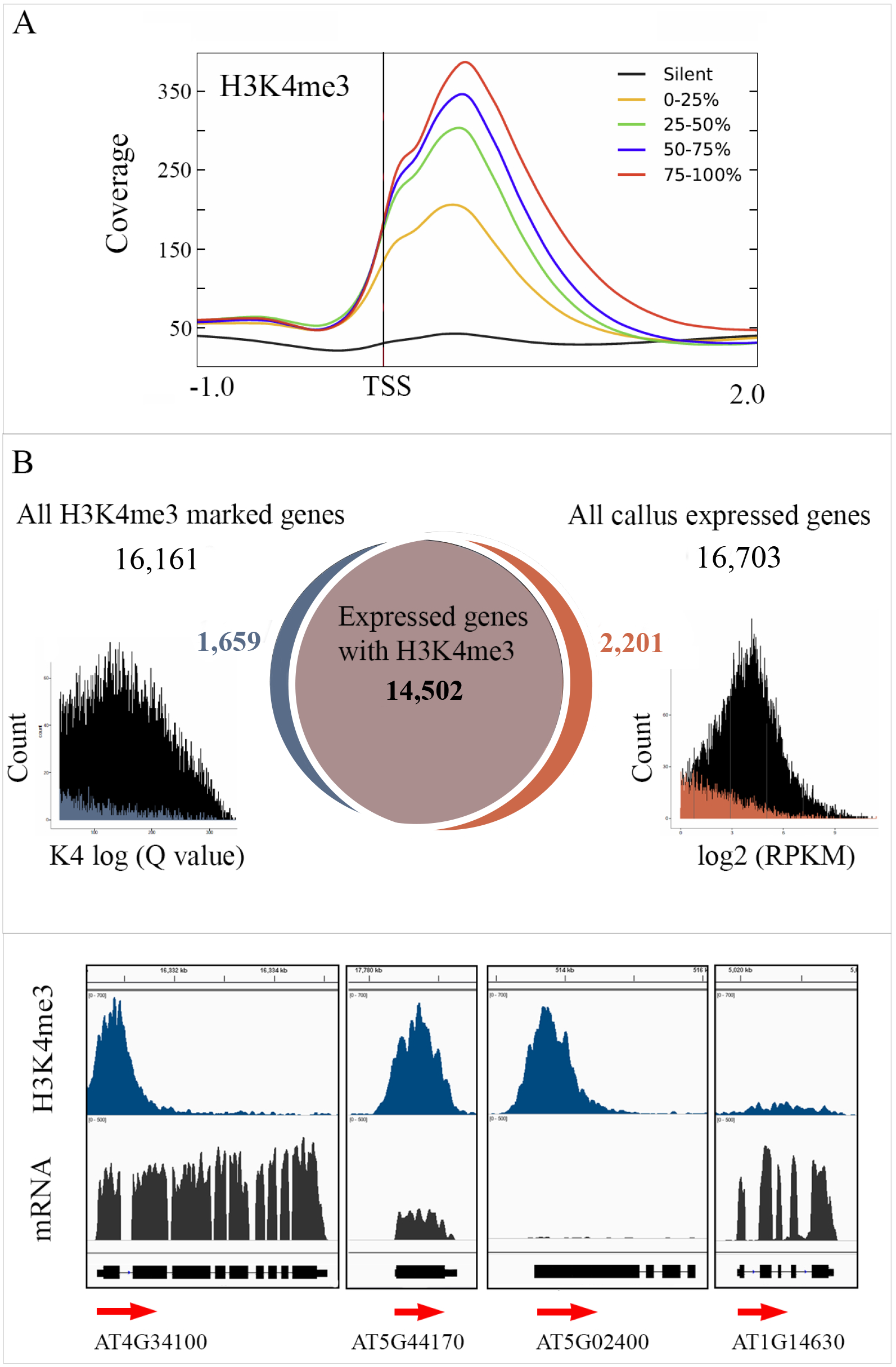
H3K4me3 correlates with gene expression. (**a**) Profile of normalized H3K4me3 signal across the TSS for genes divided into five groups: callus silent genes and four quantiles according to their expression levels. (**b**) Venn diagram showing the overlap between callus expressed genes and callus H3K4me3 marked genes. The number of genes in each category is shown. On the right a histogram of gene distribution according to their RPKM value for H3K4me3 non-marked genes (orange) and all expressed genes (black), on the left a histogram of H3K4me3 log (Q value) for H3K4me3 marked non-expressed genes (blue) and all H3K4me3 marked genes (black). (**c**) Integrated Genome Viewer (IGV) tracks of mRNA-seq and ChIP-seq signals. An example for genes with distinct correlation between H3K4me3 (blue) and expression (mRNA, black).

Profile of H3K4me3 across the TSS for all expressed genes grouped into four quantiles according to their expression levels demonstrate that H3K4me3 is highly correlated with gene activation (Fig. 2a) and therefore conserved in its association with transcription.

Overlapping the callus expressed genes with the H3K4me3 marked genes exposed three groups of genes (Fig. 2b): 14,502 expressed genes-marked by H3K4me3, which is 87% of all expressed genes, indicating that H3K4me3 serve as a general mechanism rather than regulator of specific set of genes; 2,201 expressed genes-not marked by H3K4me3, indicating that the presence of the H3K4me3 mark is not imperative for gene expression. Nevertheless, since this analysis was done on a population level, several of the genes may reflect mixed signals of H3K4me3 cooccur with transcription in some cells and neither H3K4me3 nor transcription in others. The averaging leads to H3K4me3 signal lower than the cutoff due to a dilution effect, whereas the expression level passes the cutoff; 1659 non-expressed-marked by H3K4me3 genes, that include genes like *FLOWERING LOCUS C* (*FLC*), *NGATHA3* (*NGA3*) and *BLADE ON PETIOLE 1* (*BOP1*) which were reported before to be bivalent genes, silenced and marked with both H3K4me3 and H3K27me3 (31, 32), suggesting that this group consists potential candidate for bivalency (Supplementary Data 10). Other genes in this group might be marked with H3K4me3 to be poised for later expression as was suggested for zebrafish (33). An example for genes in each group is given in Fig. 2c.

### H3K27me3 mark in callus is associated with specific set of silent genes

Mapping the H3K27me3 repressive mark in callus reveals enrichment across gene body of silenced genes (Fig. 3a), in agreement with other reports (34). However, only 3413 genes out of the 12,717 that exhibit no expression (RPKM <1) were marked with H3K27me3, indicating that this repressive mark regulates specific set of genes. Surprisingly additional 531 marked genes were found to be associated with active transcription (Fig. 3b, 3c and Supplementary Data 9), suggesting either that H3K27me3 is associated also with a distinct transcriptional outcome as reported for other organism and for Arabidopsis (35, 36), or that it is the result of averaging cells with distinct feature. Performing GO analysis on the 3413 H3K27me3 enriched and silenced genes, we identified significantly overrepresented group of genes like microRNA (43 genes out of 326), suggesting that H3K27me3 indirectly promotes gene expression by silencing miRNAs, and AGAMOUS-like (28 genes out of the 108 MADS genes in Arabidopsis). Most striking was the group of TFs, consist of 467 genes (Supplementary data 11), out of 2192 TFs in Arabidopsis (2.73-fold enrichment, p<5E-16). This group includes 202 TFs falling within the term of “developmental process”, many which are sufficient to direct cell differentiation or de novo organogenesis (Fig. 4a). One example is the set of TFs required for stomatal differentiation including *PROTODERMAL FACTOR 2* (*PDF2*), *HOMEODOMAIN GLABROUS 2* (*HDG2*) and *HDG5, MERISTEM LAYER 1* (*ATML1*), *SPCH*, MUTE and FAMA (37, 38). Another is the group of abaxial regulatory genes, which have mutual antagonistic interactions with adaxial regulatory genes to establish organ polarity. The abaxial genes are required for specifying cell identity away from the shoot apex side in all lateral organ, for example the lower side of a leaf. In the callus, genes of this group are marked by H3K27me3 and silenced (Fig.4b), consistent with the view that abaxial cell fate may be the default pattern of differentiation (39). Remarkably, the genes known to specify adaxial identity, the side close to the shoot apex, for example the leaf upper side (40), are marked by H3K4me3 and expressed in the callus.

**Figure 3.**
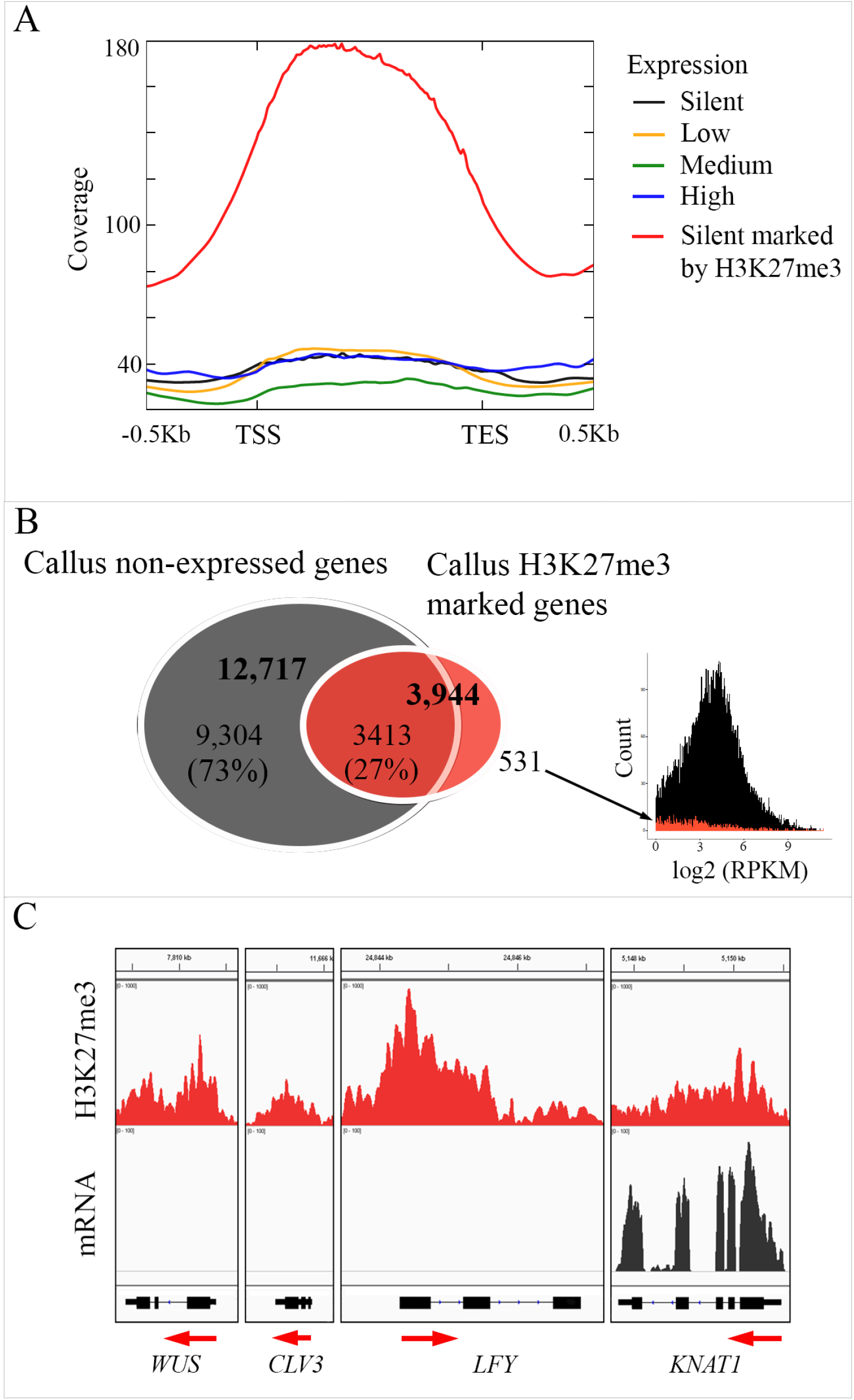
H3K27me3 mark in callus is associated with transcriptional repression of specific set of genes. (**a**) H3K27me3 exhibits broad domain of enrichment across the gene body. H3K37me3 ChIP-seq profiles across gene bodies (−0.5kb upstream to the TSS to 0.5kb downstream to the TES) for all the Arabidopsis genes divided into the following categories: silent (H3K27me3 non-marked (black) and marked (red) genes) and expressed (high, medium and low). (**b**) Venn diagram showing the overlap between callus non-expressed genes and callus H3K27me3 marked genes. The number of genes in each category is shown. On the right a histogram of gene distribution according to their RPKM value for expressed H3K27me3 marked genes (red) and all expressed genes (black). (c) Integrative Genomics Viewer (IGV) tracks of mRNA-seq and ChIP-seq signals. An example for silenced developmental genes marked by H3K27me3 and a case of exception (marked and expressed).

**Figure 4.**
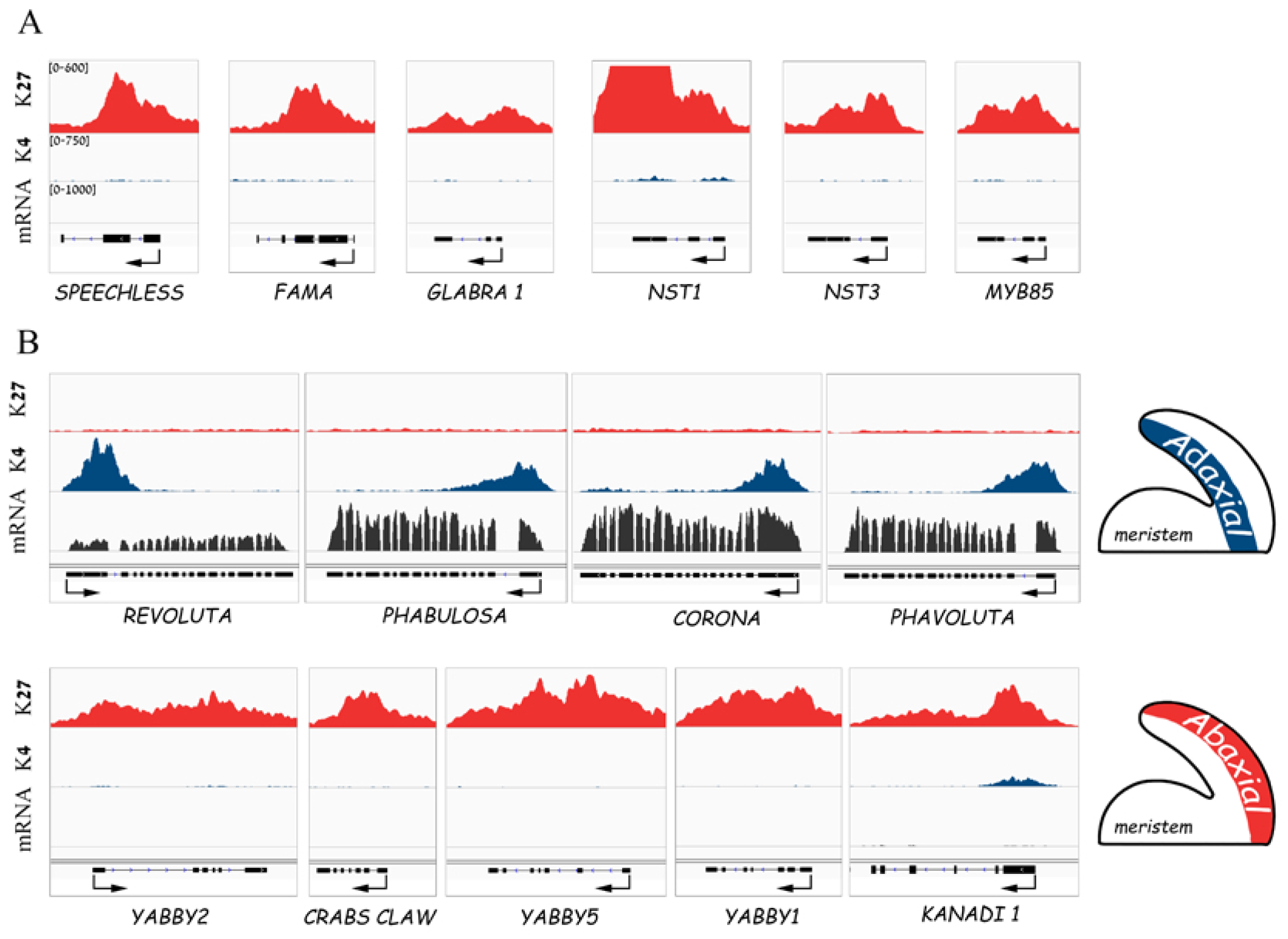
Genes sufficient to direct differentiation or de novo organogenesis are silenced in callus. IGV genome browser screenshot for mRNA and ChlP-seq data of H3K4me3 and H3K27me3. (**a**) An example for transcription factors that are sufficient to promote stomata (*SPEECHLESS* and FAMA REF), trichrome (*GLABRA1* REF) and fiber (*NST1, NST3* and *MYB85* REF) differentiation, that are marked by H3K27me3 (red) and are not expressed. (**b**) The antagonistic adaxial and abaxial identity genes display distinct H3K27me3 and H3K4me3 landscape. Upper panel: the five genes (here only four presented) specifying adaxial polarity are all marked by H3K4me3 (blue) and expressed (black) in callus. Lower panel: the *YABBY* and *KANADI* genes known to promote abaxial specification are silenced and marked by H3K27me3 (red).

From the list of 3413 genes, we also identified 339 as potentially bivalent genes that are marked with both H3K4me3 and H3K27me and show no expression (RPKM <1) Supplementary data 11). Among them we found the *FLC, NGA2, NGA3, BOP1*, and *KANADI* bivalent genes and another 44 genes that were identified as putative bivalent genes based on ChIP-seq analysis done on Arabidopsis seedlings (31), thereby confirming the quality of our analyses.

In summary, our results support the hypothesis that keeping many developmental pathways active, while maintaining dominate genes silenced to safeguard pluripotent cell identity, provides a capacity to rapidly respond to stimuli and regenerate accordingly. However, this strategy requires a mechanism to silence genes upon regenerative stimulus, to allow one developmental pathway to dominate and reinforce lineage commitment.

### *emf2-2* callus displays impaired regeneration capacity

We found that in WT callus, 467 genes encoding for TFs are marked by H3K27me3 and silenced, including at least 202 developmental regulators (Supplementary data 10). To test how prominent is the H3K27me3 mark to callus cell identity and to its capacity to regenerate, we produced callus from the PRC2 mutant *emf2-2* (hereafter *emf2*) and analyzed its capability to regenerate (Fig. 5). The *emf2* mutant was shown to be incapable of forming callus from leaf or cotyledons (7), probably due to precocious differentiation. To overcome this hindrance, we sowed the mutant directly on callus inducing media (CIM) to allow the embryonic cells to proliferate, following by cutting off the cotyledons and re-culture them on CIM. This was done for WT callus as well, and resulted in similar calli that phenotypically could not be distinguished from the *emf2* (Fig. 5a). Next, we tested the callus cell-self-identity, i.e. how the callus cells coordinate their inherent cellular programs when no external stimuli are applied. We transferred 100 calli of each genotype, WT and *emf2*, to hormone-free medium and incubated them in the dark (50 calli) or under light (50 calli) (Fig. 5b). In the dark, the WT calli developed roots that were first detected after 13 days. At day 20 all calli developed roots that kept growing even after 47 days, indicating either that with no hormonal stimuli and light signaling the root program is dominant, or that in the dark, the auxin that was absorbed on CIM is stable, and thus promote root formation. For the *emf2* calli, we observed at least one root initial on 60% of calli only from day 20, but all initials did not elongate further, and all calli started to accumulate dark brown color and eventually died (Fig. 5b).

**Figure 5.**
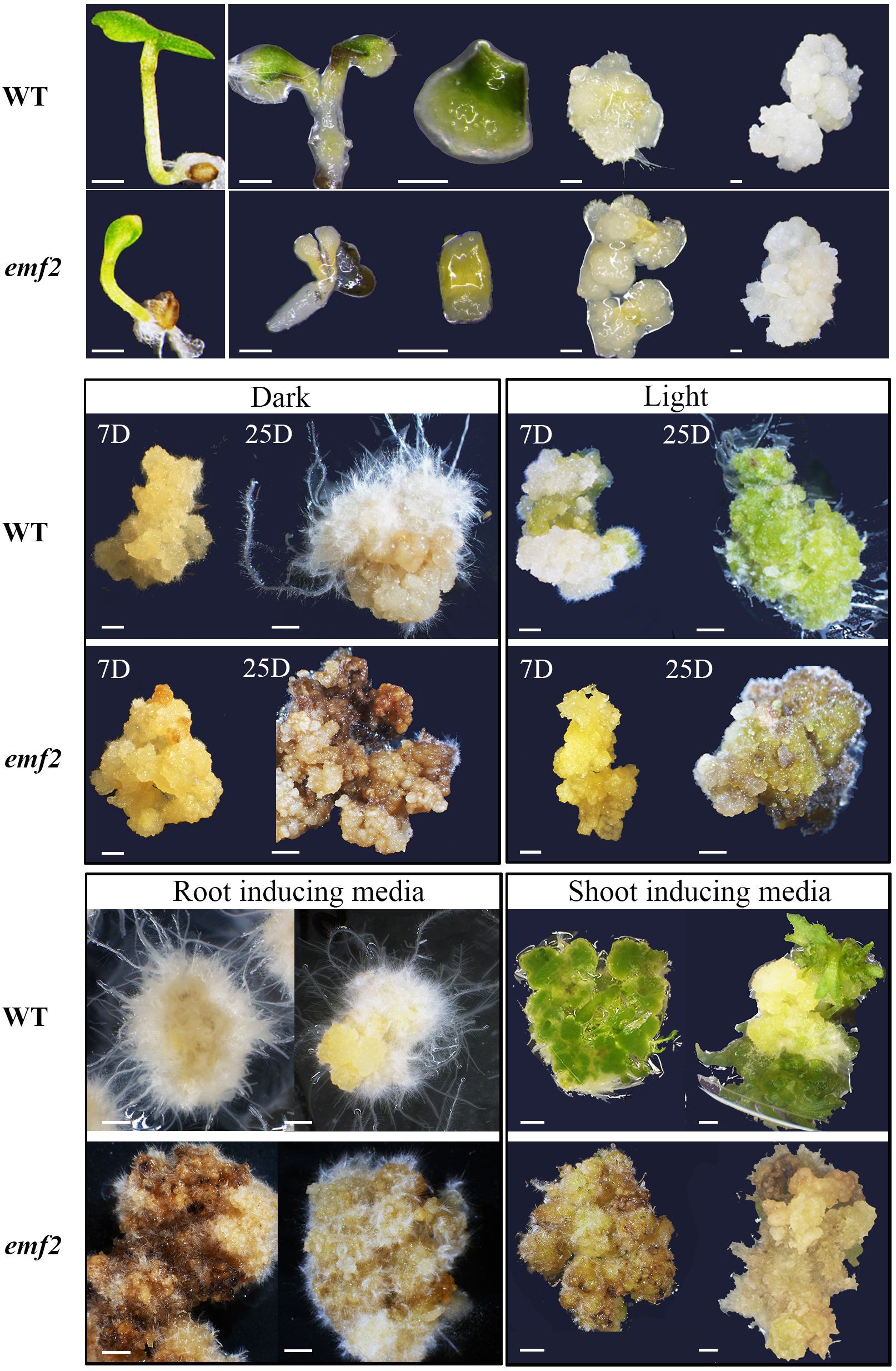
The *emf2* callus exhibits impaired regeneration capacity. (**a**) On the left, the WT and *emf2* seedlings. On the right, callus generated from cotyledons of WT and *emf2* seeds sown directly on callus inducing media (CIM) trimmed at day six and re-cultured on CIM. (**b**) Cell-self-identity test: five-week old WT or *emf2* calli were transferred to B5 media lacking hormones and cultured in dark or under light. Representative images were taken at day seven and 25. (**c**) Regeneration test: five-weeks old WT or *emf2* calli were transferred to root inducing media (RIM) or shoot inducing media (SIM). Representative images were taken at day 25 (on the left bottom side of the plate on the right upper side).

Under light, from day 7, green color accumulation appeared on 50% and 32% of WT and *emf2* calli respectively, and calli of both genotypes were still vigorous and grew in size. After 20 days, 60% and 70% of WT and *emf2* calli developed roots and root initials respectively, and from this day on the WT calli gained deep green color, while all the *emf2* calli became brown and decayed. No shoots were detected in both genotypes during the 47 days, indicating that light alone is not a sufficient stimulus to induce de novo shoot meristem formation, but it can drive chloroplast biogenesis. To test the capacity of the *emf2* callus to respond to hormonal signals and regenerate accordingly, we cultured the WT and *emf2* calli on root or shoot inducing media (RIM or SIM). For each genotype, 100 calli were cultured on RIM in the dark and 100 calli on SIM under light condition.

After 7 days on RIM, root initials started to develop on both genotypes with no significant difference along the experiment in the number of calli producing roots (Supplementary Fig X). However, the WT roots elongated and formed lateral roots, while the *emf2* roots remain short and eventually decayed in parallel to the callus dark tone accumulation. On SIM, green color accumulated on calli of both genotypes but in WT it occurred earlier, with 98% of the calli displayed green sections already on day four as compared to 42% in *emf2*. First shoots were observed in both genotypes on day 11. However, on day 15, 96% of the WT calli regenerated several vegetative shoots on each callus, while only 8% of the *emf2* calli regenerated single shoot at the reproductive phase (Supplementary Fig X). Unlike on RIM plates, on day 25 the *emf2* calli did not show signs of cell death and decaying at this stage, but after 30 days all calli on SIM as on RIM decayed, while the WT calli were still vigorous (Supplementary Fig X).

Taken together these results indicate that in the absence of functional EMF2 complex the callus cells fail to acquire new cell identity.

### *emf2* callus displays minor changes in gene expression

Seedlings of *emf2* were shown to display reduced H3K27me3 tri-methylation on 54% of the WT marked genes (41). To study how mutation in the *EMF2* gene affects transcription in callus, we performed differential gene expression analysis on mRNA-Seq from WT and *emf2* calli. This analysis revealed unexpected results. In total 16,646 genes were expressed in *emf* callus (RPKM>1, Supplementary data 12) with only 812 genes exhibiting differential expression (p<0.05): 374 up- and 438 down-regulated genes in *emf2* calli as compared with WT, leaving 15,834 genes with similar expression levels (Fig. 6a Supplementary data 13). This low numbers are surprising based on the analysis done on WT and *emf2* seedlings, revealing more than 2,500 genes that are differentially expressed (41), nevertheless the seedlings from the two genotypes composed of different tissues, vegetative in WT and reproductive in *emf2*. The *VRN2* gene, that like the *EMF2* encodes for a protein with homology to Drosophila *Su(z)12* (42) and serves as a PcG components (43) exhibits high expression level (RPKM =7), in both WT and *emf2* calli (Table 1), suggesting that functional redundancy might compensated for the loss of EMF2 function.

**Figure 6.**
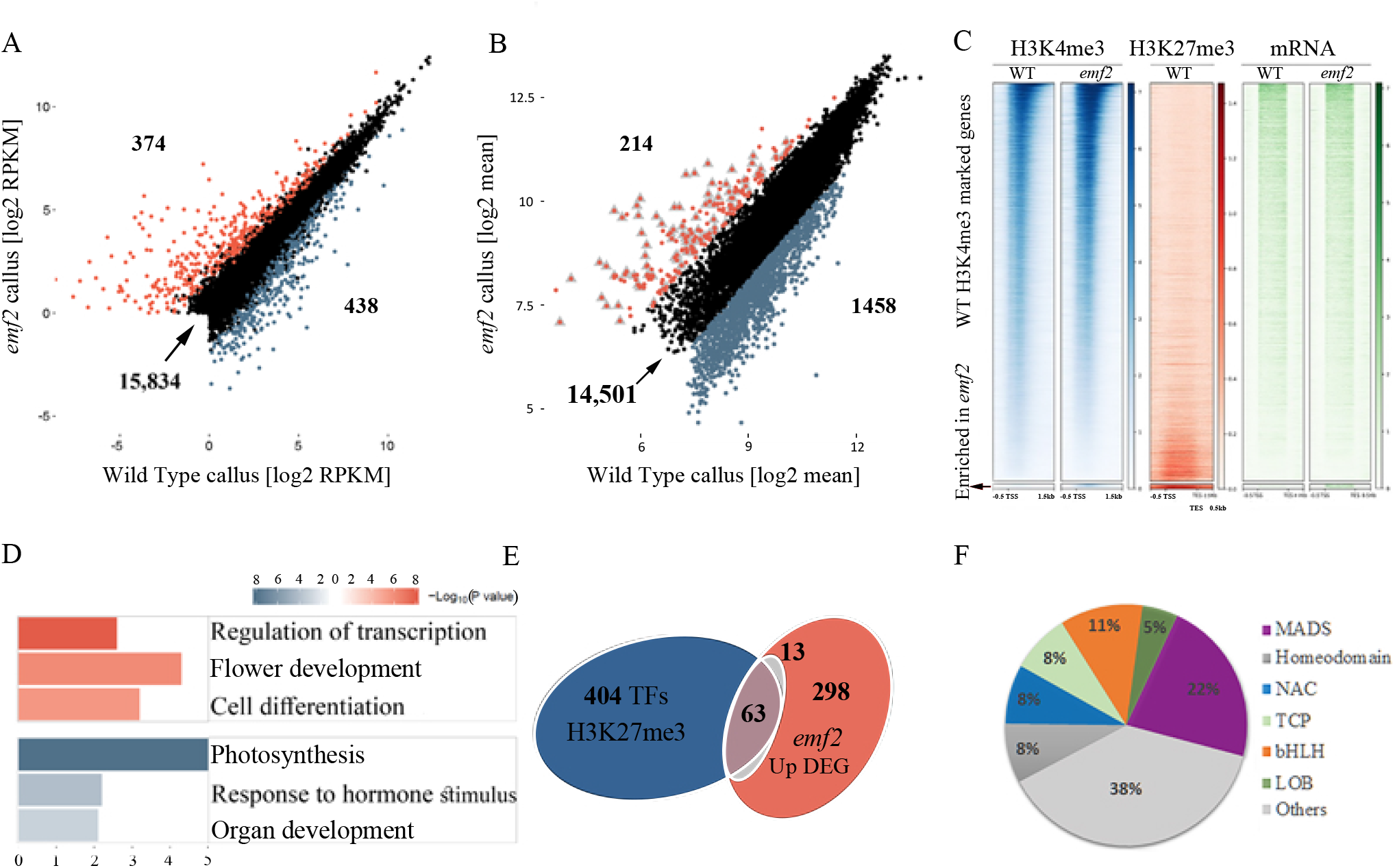
The *emf2* callus exhibits minor alterations in gene expression and H3K4me3 acquisition with enrichment in up-regulated genes coding for transcription factor. (**a**) differential expression between *emf2* and WT callus from RNA-Seq normalized values. Scatter plot demonstrates mean expression vs. log2 fold-changes in expression. Red and blue dots indicate *emf2* significant up or down-regulated genes (P < 0.05). (**b**) Differential H3K4me3 modifications between *emf2* and WT calli presented as scattered plot of significant enrichment (red) and reduction of the mark. Gray triangles mark the 94 genes that are marked by H3K27me3 in WT. (**c**) H3K4me3 displays anticorrelation with H3K27me3 and strong correlation with expression. Heat maps for WT H3K4me3 marked genes only, sorted by the H3K4me3 signal form the highest to lowest. Maps present the pattern for H3K4me3 and H3K27me3 ChIP-seq occupancy −0.5kb from TSS and 1.5kb (K4) or 5kb (K27) downstream to TSS and mRNA-seq for WT and *emf2* callus. The box at the bottom represent X genes that acquired H3K4me3 mark in the *emf2* callus (showing no H3K4me3 signal in WT, strong H3K27me3 signal in WT high expression in *emf2*). (**d**) Euler diagram showing significant enrichment (p-value of 5.32e-16) in genes encoding for transcription factors (76, yellow) among the *emf2* up-regulated expressed genes (374, red) The P-value was calculated by a hypergeometric test using the following numbers: 2192 TF out of 29420 genes in Arabidopsis (according to TAIR) and 76 TF out of the 374 *emf2* up-regulated genes. The number of genes in each category is shown. (**e**) **Motives or specific GO**

Out of the 374 *emf2* callus up-regulated genes, 131 are silenced (RPKM<1) in WT. To further investigate how the differentially expressed genes might affect the capacity to regenerate we performed GO analysis (Fig. 6d, Supplementary data 14). The most enriched biological processes category for the 374 up-regulated genes was the “transcription factor activity” term, as 76 genes encode for TFs, from which 63 were marked by H3K27me3 in WT callus (supplementary data 15). Out of the 63 TFs, 14 genes belong to Type-II MICKC sub-family of the MADS-box TFs, that control flowering transition and formation (44). For example, the *SEPALLATA 2* gene required for petals, stamens and carpels development (45), is marked with H3K27me3 in WT callus and shows no expression (RPKM of 0.06), whereas in *emf2* exhibits high expression level (RPKM of 8.5, Log2 Fold Change =7). The *AGAMOUS-LIKE 72* (*AGL72*) TF, that involved in floral transition (46), shows RPKM value of 9.4 in *emf2* and no expression in WT (|log2 Fold Change = 8.5). This is also reflected in the significant enrichment of the “flower development” term (GO:0009908, p< 8.E-07) and is consistent with the *emf2* phenotype of flowering upon germination (47), and the regeneration of flower from the *emf2* callus (supplementary Fig X). Remarkably, 10 up-regulated genes in the *emf2* calli are associated specifically with carpel development (48). For example, *SEEDSTICK* (*STK*), that was shown to be sufficient to induce the transformation of sepals into carpeloid organs and to promote carpel development in the absence of AG activity (49), showed Log2 Fold Change of 7.4. What caught our attention is the enriched GO term “cell differentiation: (GO:00030154, p< 8E-08), since the expression of genes that promote cell differentiation can contribute to the reduced capacity of the *emf2* callus to regenerate. Up-regulated genes in this term include TFs involved in xylem fibers differentiation like *VASCULAR-RELATED NAC-DOMAIN 3* and *6* and *ACAULIS 5* (50), cuticle synthesis (*CUTICULAR 1*), hair cell differentiation (*HDA18*) and leaf cellular differentiation (*TCP* 2, 10 and 17) (51), suggesting that EMF2 represses those genes in WT callus to prevent cell differentiation, thereby safeguard pluripotency identity.

The *emf2* callus also exhibit 438 down-regulated genes and the decrease in expression level might be due to indirect effect. For example, the decrease can be the outcome of up-regulation of 76 genes encoding for TFs, that might initiate distinct gene regulatory networks, or the result of up-regulation of 43 microRNA genes. To test this, we preformed GO analysis on the 438 down-regulated genes and revealed remarkable enrichment in categories related to photosynthesis (Supplemented data 14), consistent with previous report showing similar result with *emf2* seedlings (52). For example, 33 out of the 113 genes in the biological processes term Photosynthesis (GO:0015979, p< E-31), and 48 genes out of 322 in the cellular component term Thylakoid (GO:0009579 p< E-31) are down regulated genes.

### *emf2* exhibits minor changes in H3K4me3

The H3K4 histone methyltransferases are key players in counteracting PcG-mediated repression (4), but how PcGs affect their activity is still not clear. To study the effect of the mutation in *EMF2* on the H3K4me3 distribution, we performed ChIP-seq analysis on *emf2* callus using H3K4me3 specific antibodies, and compared it with WT callus (Fig. 6b). In *emf2* callus, the H3K4me3 mark was detected on 16,173 of the genes (Supplementary Data 14), from which 90% (14,562) are expressed (RPKM>1), demonstrating significant positive correlation with expression similarly to WT (Fig 2).

We next analyzed the differential levels of enriched H3K4me3 peaks between WT and *emf2* calli using edgeR algorithm (53) (Supplementary data 15). Surprisingly only 214 genes exhibited significant greater signals in *emf2* callus (|log2 Fold Change| > 1 and FDR < 0.01), from which 109 genes are marked by H3K27me3 in WT, including 35 genes encoding for TFs, most of which involved in plant development (Fig 6f).

Most of the H3K4me3 marked genes showed no significant differences in the signal intensity between the two genotypes (Fig 6b), and strikingly, 1458 genes had weaker signal in the *emf2* callus as compared with WT. Out of the 1458 genes, 726 exhibit no expression in both genotypes, regardless of the present of H3K4me3 and only 333 were marked by H3K27me3 in WT, suggesting that the reduced H3K4me3 signal in this group of genes is indirect.

To summarize our analyses, we generated heat maps for WT H3K4me3 marked genes only, sorted by the H3K4me3 signal form the highest to lowest, demonstrating that H3K4me3 displays anti-correlation with H3K27me3 and strong correlation with expression (Fig 6c). The *emf2* callus exhibits similar pattern except for the 56 genes that are marked by H3K27me3 in WT, acquired the H3K4me3 mark in the *emf2* and gained expression (The box at the bottom of the heat maps and Fig 7).

**Figure 7.**
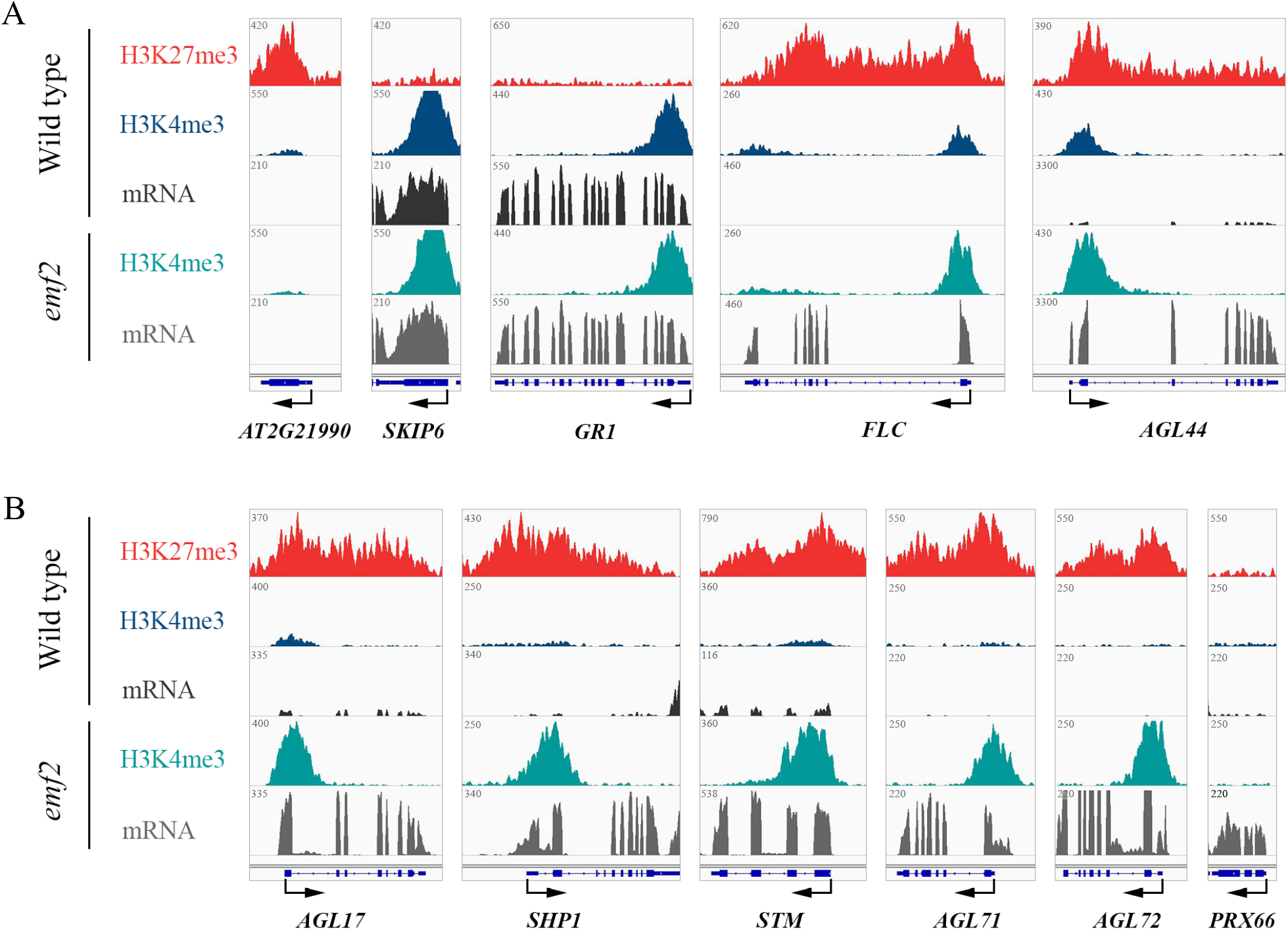
H3K27me3, H3K4me3 and mRNA IGV tracks for selected genes. (**a**) a gene that in WT is marked by H3K27me3 (red) and shows no enrichment in H3K4me3 and no expression both in WT and in *emf2* (left); genes that in WT show no enrichment for H3K27me3 and are marked by H3K4me3 and are expressed both in WT and *emf2* (middle); genes that in WT are marked by both H3K27me3 and H3K4me3 and show no expression (potential bivalent genes) and in *emf2* show enrichment in H3K4me3 and are expressed (right). (**b**) Example for genes that gained expression in the *emf2* callus. Genes that in WT are marked by H3K27me3 and show no H3K4me3 enrichment or expression and in *emf2* acquired H3K4me3 and are expressed. On the right and example for genes that gained expression without H3K4me3. Units are of total cpm (count per million).

## Discussion

The question of how animal stem cells maintain their pluripotent identity is the subject of intensive study. In contrast, much less is known on similar systems in plants. In this work we provide insights into callus features, having specific gene expression patterns that sustain high proliferation rate, independently from maintaining the potential to initiate all lineages of the mature organism. It is generally accepted that mammalian embryonic stem (ES) cells maintain pluripotency through retaining globally open chromatin state by basal transcription, to confer transcriptional competence for developmental programs (54, 55). We identify related strategy in callus, of keeping lineage-affiliated genes transcriptionally active, enabling the rapid response to signals, without the needs to go through the complex process of transcription. However, such strategy requires complementary mechanism to prevent precocious differentiation. We propose two independent mechanisms for maintaining callus identity; one of keeping the cell cycle machinery active, which impact cell fate decisions (56). This is supported by the observation that upon culturing the callus on media devoid of proliferation instructive signals, the cells differentiated (Fig. 5). Contrarily to the highly proliferative characteristic of callus cells, central feature of plant pluripotent stem cells is the slow mitotic rate (18, 57, 58), demonstrating that pluripotency it not necessarily a characteristic unique to slow dividing cells; another mechanism, is the silencing of dominant TFs via epigenetics modifications that are maintained thorough constant proliferation.

Pluripotent cells primarily possess dual capacities: maintaining cellular identity of less differentiated cells and having the capacity to form multiple cell lineages. This requires plasticity to reprogram the epigenetic states upon signals, to establish and maintain the lineage-specific transcriptional program and to shut down other programs. Here we show that tri-methylation of H3K27 plays a major role in maintaining callus cellular identity and in providing the required plasticity. We demonstrated that the *emf2* callus failed to orchestrating proper differentiation and regeneration, indicating that in the absence of deposition of the trimethylation on H3K27 on specific genes the capacity to form multiple cell lineages is impaired.

We show that *emf2* callus can be established and proliferate in culture, indicating that hormone perception and signal transduction are not compromised. In addition, it indicates that the EMF2 component of the PRC2 is dispensable for cell proliferation, consistent with the view that self-renewal in mammalian ES cell is generally not subjected to repressive epigenetic mechanisms (59). We found that only 812 genes are differentially expressed between *emf2* and WT. For the insignificant number of 374 up-regulated genes in *emf2*, we can propose several scenarios and explanations: 1. EMF2 might regulate small set of genes. However previous study showed 54% reduction in total H3K27me3 in *emf2* seedlings compared with WT (41); 2. VRN2, which has similar role to the EMF2 in the PRC2 complex and is highly expressed in both WT and *emf2* calli, targets overlapping genes and largely compensate for the loss of EMF2 function. That scenario highlights the exact set of gene specially regulated by EMF2: marked by H3K27me3 in WT callus and gained the H3K4me3 mark and expression in *emf2;* 3. There is a massive reduction in H3K27me3 in *emf2*, but the removal of a repressive mark does not necessarily activate genes. This was demonstrated for example, in mouse ES cells carrying a mutation in *Ezh2*, the HMTase component of PRC2, where the loss of H3K27me3 mark in numerous loci resulted in transcriptional activation of only one third of the marked loci (60).

The normal development of *emf2* callus, despite the up-regulation of 76 genes encoding for TF, from which 63 are marked by H3K27me3 and silenced in WT callus, some of which can promote differentiation and organogenesis, suggests that the signal for keeping the cell cycle machinery active is more robust. Yet, the failure to differentiate upon removal of the proliferative signal or in response to regenerative signals, indicates that acquisition of new cell-fate requires epigenetic specificity, and suggests that activating lineage-specific gene networks to commit to a single developmental path requires a capacity to turn off pluripotency networks.

Several MADS-box TFs were shown to be involve in downregulating the expression of photosynthetic genes to confer flower identity (61, 62). Therefore, the enrichment in photosynthesis related gene among the *emf2* callus-downregulated genes, might be the direct effect of *MADS*-box genes up-regulation in *emf2*. The activation of the *LOB domain 29* (*LBD29*) TF in the *emf2* callus, which was shown to directly repress genes involved in photosynthesis (63), might also contribute to this enrichment.

In Arabidopsis, mutations in *EMF2* induce immediate flowering after germination (64). In general, flowering can be promoted by environmental cues such as long-day photoperiod or low-temperature, leading to the activation of floral integrators genes, *FLOWERING LOCUS T* (*FT*), *SOC1, TWIN SISTER OF FT* (*TSF*) and *LFY* (65). Those TFs induce the transition to flowering, leading to floral meristem specification, followed by the activation of a downstream genetic networks comprises mainly of *MADS*-box genes, to specify floral organ identities (66). In our study we cultured the callus at conditions that do not promote flowering (under dark at 22C^0^), and the four integrators genes showed no expression in both WT and *emf2*. However, many of the downstream MADS-box genes are up-regulated in *emf2*, demonstrating that activation of floral organ specification genes is independent of the transition to flowering. This result suggests that the four integrators are required solely to release the suppression and not for direct activation of the downstream MADS-box genes.

No strong consensus has been reached on the level of callus homogeneity, and the question of whether the callus cells have similar cellular identity has yet to be investigated (1). Our comparative analysis between WT and *emf2* calli identified only 324 up-regulated genes, demonstrating the similarity of the two calli, suggesting that the callus produced form embryonic tissue is more unify. This can be tested in the future by performing a transcriptomic analysis at single-cell resolution (67), with the recognition that it is not a bias-free method.

Our approach offers an excellent solution to yield unbiased results for functional genomic studies and the study of pluripotency or for transcriptome and chromatin state comparison between various mutants that do not have other comparable tissues or organs. Using our experimental system, we were able to pinpoint the potential genes regulated solely by EMF2, mainly floral organ specification genes. Future challenge of performing ChIP-seq analysis for EMF2 can yield a data on all the EMF2 direct targets.

In summary, we provide insights into H3K27me3 function in establishing pluripotent cell identity in callus, by silencing key TFs and contributing to the capacity to initiate new cell lineage. Our findings also provide the starting point for further studies on the genome-wide dynamics of H3K4me3 and H3K27me3 taking place during regeneration.

## Methods

### Plant materials and growth conditions

Arabidopsis thaliana accession Columbia-0 (Col-0) was used in this work as wild type and a mutant of the *EMF2* gene (AT5G51230) the *emf2-1* mutant (ABRC germplasm CS16238) on Col-0 background (47). The mutant was genotyped according to Calonje et al. 2008 (68). For all experiments, seed were surface sterilized for 10 minutes in 3% Sodium Hypochlorite containing 0.1% Triton X-100, washed 4 times with DDW and sown on Medium containing Murashige and Skoog (MS) basal salt mixture. Plates were placed at 4^0^C for 2 day in the dark. and then transferred to a continuous light growth room at 23^0^ C for one week. Next, seedlings were transplanted to soil and grown in long day (16h light/ 8h dark) at 21C. Rosette leaves were harvested 2 weeks after transplantation.

### Tissue culture

All tissue culture experiments were done in a growth room at 23^0^C under continuous light or dark conditions. To generate callus from Col-0 and *emf2-1* mutant cotyledons, sterilized seeds were sown directly on callus inducing medium (CIM) (Gamborg B5 medium with 0.5gr/L MES and supplemented with 2% dextrose, 0.9% phytagel, supplemented with 2.2μM 2,4-dichlorophenoxyacetic acid 2,4-D and 0.46μM kinetin. Following two days of incubation at 4^0^C in the dark, plates were transferred to continuous light growth room. After seven days, cotyledons with callus like formation were trimmed and transferred to a new CIM plate placed in dark. During the next 5-6 weeks, calli were re-cultured to new CIM plates every 7-10 days. All analyses were done on 6-8 weeks old calli.

### Callus internal identity experiments

Calli were transferred to Gamborg B5 medium deprived of hormones, under continuous light or dark conditions. Calli were monitored under a stereomicroscope and photographed

### Competency to regenerate tests

Calli of WT or *emf2-1* mutant and 21 days old WT rosette leaves were transferred to either Shoot Inducing Media (SIM): Gamborg B5 medium supplemented with 4.4μM 2-isopentenyladenine (2iP) and 0.5μM 1-Naphthaleneacetic acid (NAA) under continuous light conditions or Root Inducing Media (RIM): Gamborg B5 medium supplemented with 0.5μM NAA under dark conditions. Calli and leaves were monitored under a stereomicroscope and photographed.

### RNA extraction

In order to correlate chromatin characteristics with gene expression, tissues from each ChIP experiment were fast frozen in liquid nitrogen. Samples were grinded and total RNA was extracted using the RNeasy Plant Mini Kit (Qiagen) following the manufacturers instructions Contaminating genomic DNA was removed from the RNA samples by treatment with RNase-free DNase (Qiagen). Each RNA sample was loaded on agarose gel for integrity validation and quantified using NanoDrop.

### RNA-Sequencing

Samples were processed for sequencing at the Thechnion Genome Centre (TGC) according to Illumina’s protocols. TruSeq RNA V2 sample prep kit (RS-122) was used. Total RNA was polyA-selected followed by fragmentation and random hexamer primed reverse transcription. After second strand synthesis, indexed adapters were added and cDNA was amplified by PCR during 15 cycles. Every 4 indexed libraries were loaded on to an individual lane of the Illumina’s HiSeq 2500 system.

### Chromatin immunoprecipitation

ChIP was performed on nuclei isolated from callus as described (69) with modifications to adjust for callus. Detailed protocol is found in Supplementary materiel and method file. In short, Six-week old calli samples were harvested from 6 plates and pooled into 4 tubes of 1-1.5gr and were subjected to crosslinking.

### ChIP validation

For validation of the ChIP experiment, a comparative semi-quantitative PCR between positive and negative DNA binding sites was performed. For each factor, two sets of primer were design: positive control (PC), for estimated positive DNA binding site and negative control (NC), for estimated negative binding site (Table 1 Supplementary materiel and method file). The two sets of primers were tested on Input DNA sample diluted 1:1,000 and IP sample. The reaction mix included: 7.5μl of GoTaq Green master mix (Promega), 0.4μl of each primer (10μM) and 5.9μl diluted DNA. The PCR program started at 95^0^C for 3min followed by 36-40 cycles of 95C^0^ for 30s, annealing temperature (calibrated to each set of 4 primers) for 30sec and 72^0^C for 30sec. Along the PCR reaction, samples were taken out from cycle 28 and every 2 cycles and loaded on 2% agarose gel.

### Chromatin immunoprecipitation sequencing

Samples were processed for sequencing at the Thechnion Genome Centre (TGC) according to Illumina’s protocols.

The ChIP-seq libraries were prepared using, TruSeq Nano DNA Library Prep Kit (FC-121) was used. ChIP enriched DNA fragments were size selected on an agarose gel to enrich for fragments 200bp in size, indexed adaptors (diluted 1/10) were added and the library was amplified using PCR during 14 cycles. Every 7-10 indexed libraries were loaded on to an individual lane of the Illumina’s HiSeq 2500 systems for collecting 50 base-pair single-end.

Total reads count and unique reads count are presented in table 2 (Supplementary).

### Next Generation Sequencing (NGS) analysis

Each NGS analysis included quality control analysis using FastQC (70) and cutting off the reads adaptors using cutadapt (71) NGS reads were mapped to Arabidopsis thaliana reference transcriptome TAIR10 using TopHat 2 (Kim 2013) for RNA-Seq and Bowtie2 (72) for ChIP-Seq. Integrative Genomics Viewer (IGV) was used for the visualization of the data (73, 74). Biological replications consistency was measured using spearman correlation of BAM files in deepTools (75). RNA-Seq samples with p>0.96 and ChIP-Seq samples with p>0.88 were merged.

## Supporting information

Sup Materiel and Methods

