## Supplementary material for "The H3K27me3 epigenetic mark is crucial for callus cell identity and for the acquisition of new fate during root and shoot regeneration": Sup Materiel and Methods

**Chromatin immunoprecipitation**

ChIP was performed on nuclei isolated from callus as described (Kaufmann et al. 2010a) with modifications to adjust for callus. Detailed protocol is found in Supplementary materiel and method file.

**Tissue collection:** Six-week old calli samples were harvested from 6 plates and pooled into 4 tubes of 1-1.5gr (4-6gr in total). For leaf: 21 days old rosette leaves were harvested from about 400 plants and pooled into 8 tubes of 1-1.5gr (8-12gr in total). After all tissue was collected, 0.5gr was fast freeze in liquid nitrogen for RNA extraction. The rest was subjected to crosslinking.

**Crosslinking:** Collected tissue was immersed in 35ml ice cold MC buffer (10mM sodium phosphate pH=7, 50mM NaCl and 100mM sucrose) containing 1% Formaldehyde. Calli samples were fixated for 10min, while leaves samples were fixated 3 times each for 5min under vacuum (700mmHg). To stop the crosslinking, 3.5ml of glycine 1.25M was added to the solution for 5min under vacuum for leaves and with no vacuum to calli. Next, samples were washed with 200ml MC buffer on a sieve, carefully dried on KimWipe paper, transferred to 50ml falcon and fast freeze in liquid nitrogen.

**Nuclei enrichment:** Tissue was grinded using cold mortar and pestle, transferred to 20ml M1 buffer (10mM sodium phosphate pH=7, 100mM NaCl, 1M Hexilene-glycol, 10mM 2-mercaptoethanol and complete protease inhibitor cocktail) and shacked slowly to allow all powder to melt in the solution. Solution was filtered through doubled layered Miracloth (Calbiochem) soaked in M1 buffer to a new 50ml falcon and centrifuged at 2500g for 20min at $4^0$C. Supernatant was carefully discarded. Pellet was re-suspended in 10ml of M2 buffer (10mM sodium phosphate pH=7, 100mM NaCl, 1M Hexilene-glycol, 10mM $MgCl_2$, 0.5% Triton X-100, 10mM 2-mercaptoethanol and complete protease inhibitor cocktail) and shacked slowly on ice for 10min, followed by 10min centrifugation at 2500g in $4^0$C. After supernatant was carefully discarded, pellet was washed once again in 10ml M2 buffer and one more time in 10ml M3 buffer (10mM sodium phosphate pH=7, 100mM NaCl, 10mM 2-mercaptoethanol and complete protease inhibitor cocktail).

**Chromatin sonication:** To each tube of isolated nuclei , 0.5ml of sonic buffer (10mM sodium phosphate pH=7, 100mM NaCl, 0.5% Sarkosyl and 10mM EDTA) was added and four or eight tubes (for calli or leaves samples respectively) were pooled to one sample in 15ml tube. From each sample, 100µl for +DC control and 100µl for –DC control were set aside ($4^0$C) and the rest was aliquoted to 450-500µl in 1.5ml tubes. Samples were sonicated using the cup horn device in S-4000 sonicator by MISONIX. In order to achieve smear range of 100-700bp, sonication program was as followed: Amplitude=2, ON time=60sec, OFF time=30sec, pulse number=15. Following sonication, tubes were centrifuged at 13,000g for 10min at $4^0$C and supernatants were transferred to new tubes. Second centrifugation was carried and clear supernatants were joined together to a 15ml tube to which equal amount of IP buffer (50mM HEPES pH=7.5, 150mM NaCl, 5mM $MgCl_2$, 10µM $ZnSO_4$, 1% Triton X-100, and 0.5% SDS) was added. 250µl from the sample were set aside ($4^0$C) to serve as Input (In) control.

**Immunoprecipitation**

100-150µl (for 4 or 8 starting tissue tubes respectively) of protein A or G magnetic beads were washed twice with 1ml IP buffer in LoBind tubes (Eppendorf). 1ml IP buffer and relevant antibody (according to table 3) were added to the washed beads and incubated for 4-5h in slow rotation at $4^0$C to induce beads-antibody complexes. After 5 washes with 1ml IP buffer, complexes were added to the sonicated chromatin sample (in 15ml tube) for overnight incubation under slow rotation at $4^0$C.

**De-crosslinking**

Next day, beads-antibody-protein-DNA complexes were washed 4 times with 1ml IP buffer. Finally, 0.5ml IP buffer was added. IP buffer was also added to all controls (+DC, -DC and Input) to reach 0.5ml. To all samples, 5µl of RNase A (1mg/ml, Sigma) were added and samples were incubated at 37C for 2-3h. After –DC control sample was set aside ($4^0$C), 15µl of Proteinase K (20mg/ml, Roche) were added to the samples for overnight incubation at 37C.

**DNA isolation**

Next day, following De-crosslinking of the beads-antibody-protein-DNA, beads were discarded and all samples were subjected to phenol/chloroform DNA extraction followed by ethanol precipitation. Because of low DNA recovery from the IP sample, 2µl of glycogen were added to the IP sample at the precipitation step. Finally, 30µl DDW were added and DNA concentration was measured (not IP sample) using NanoDrop machine.

**ChIP validation**

For validation of the ChIP experiment, a comparative semi-quantitative PCR between positive and negative DNA binding sites was performed. For each factor, two sets of primer were design manually or using the Primer3Plus (http://www.bioinformatics.nl/cgi-bin/primer3plus/primer3plus.cgi). A set for positive control (PC), design for estimated positive DNA binding site of the factor, and a set for negative control (NC), for estimated negative binding site (**table 2**). The two sets of primers were initially tested on Input DNA sample diluted 1:1,000. The reaction mix included: 7.5µl of GoTaq Green master mix (Promega), 0.4µl of each primer (10µM) and 5.9µl diluted DNA. The PCR program started at $95^0$C for 3min followed by 36-40 cycles of 95C$^0$ for 30s, annealing temperature (calibrated to each set of 4 primers) for 30sec and 72$^0$C for 30sec. Along the PCR reaction, tubes were taken out from cycle 28 and every 2 cycles and loaded on 2% agarose gel. If both bands (PC+NC) showed similar strength along the elevating cycles, the same reaction will be done again including the IP sample. However, because IP DNA is very limited, it was used only for one PCR reaction at the most elevated cycle chosen according to the Input calibration. Again, PCR products were loaded on 2% agarose gel and bands were analyzed for enrichment.

***Table 2. List of primers used in this work***

| **Primer Name** | **Sequence** | **comment** |
|---|---|---|
| | **Genotyping** | |
| *emf2-1*geno_fwd | AACAACAAATTGCAGAAG | *emf2-1* genotyping (427) |
| *emf2-1*geno_rev | CTTGGATATCATTGTCTCA | *emf2-1* genotyping (428) |
| | **ChIP validation** | |
| SAG12_first_exon_fwd | AAGGAGGAAAACAATCGCTAC | *Negative for H3K4 leaf and *emf2-1* calli<br>*Positive for H3K27 calli and leaf ([illegible]) |
| SAG12_first_exon_rev | GCAAACTGATTTACCGCAAG | *Negative for H3K4 leaf and *emf2-1* calli<br>*Positive for H3K27 calli and leaf ([illegible]) |
| SAG12_downstream_fwd | CGTCATCGTCTTCTTCTTTCTC | *Positive for H3K4 leaf and *emf2-1* calli<br>*Negative for H3K27 leaf ([illegible]) |
| SAG12_downstream_rev | GCTATGCCAGAAACCTTCGT | *Positive for H3K4 leaf and *emf2-1* calli<br>*Negative for H3K27 leaf [illegible] |
| ACT2_fwd | GCTGGATCTCGATCTTGTTTTC | *Negative for H3K27 calli (444) |
| ACT2_rev | CTCTCCATCAAGGTCAAGCC | *Negative for H3K27 calli (446) |
